# Subjugation of TGFβ Signaling by Human Papilloma Virus in Head and Neck Squamous Cell Carcinoma Shifts DNA Repair from Homologous Recombination to Alternative End-Joining

**DOI:** 10.1101/353441

**Authors:** Qi Liu, Lin Ma, Trevor Jones, Luis Palomero, Miquel A. Pujana, Haydeliz Martinez-Ruiz, Patrick Ha, John Murnane, Isabel Cuartas, Joan Seoane, Michael Baumann, Annett Linge, Mary Helen Barcellos-Hoff

**Author notes:** Biology of Human Tumors. This study was funded by the UCSF Department of Radiation Oncology to MHBH. **Corresponding Author: Mary Helen Barcellos-Hoff, Ph.D.** Professor and Vice Chair Department of Radiation Oncology University of California, San Francisco 2340 Sutter Street San Francisco, CA 94143. The prognosis for oropharyngeal HNSCC patients suggests that human papilloma virus (HPV) confers vulnerability to standard-of-care radiation and chemotherapy. Although HPV impairs p53 and retinoblastoma proteins, it can also compromise TGFβ signaling. We show that loss of TGFβ signaling leads to homologous recombination deficiency that increases sensitivity to radiation and chemotherapy, but the penultimate basis for poor DNA damage repair is the shift to error-prone, alternative end-joining repair (Alt-EJ) requiring both PARP1 and POLQ. The loss of TGFβ signaling in HPV-positive HNSCC is an experiment of nature that underscores a novel route by which TGFβ inhibitors can be exploited clinically in poor prognosis HPV-negative HNSCC.

## Abstract

**Purpose:** Following cytotoxic therapy, 70% of patients with human papillomavirus (HPV) positive oropharyngeal head and neck squamous cell carcinoma (HNSCC) are alive at 5 years compared to 30% of those with similar HPV-negative cancer, which is thought to be due to dysregulation of DNA repair. Loss of transforming growth factor β (TGFβ) signaling is a poorly studied consequence of HPV that could contribute to this phenotype.

**Experimental Design:** Human HNSCC cell lines (n=9), patient-derived xenografts (n=9), tissue microarray (n=194), TCGA expression data and primary tumor specimens (n=10) were used to define the relationship between TGFβ competency, response to DNA damage, and type of DNA repair.

**Results:** Analysis of HNSCC specimens *in situ* and *in vitro* showed that HPV associates with loss of TGFβ signaling that increases the response to radiation or cisplatin. TGFβ suppressed miR-182 that inhibited both BRCA1, necessary for homologous recombination repair, and FOXO3, which is required for ATM kinase activity. TGFβ signaling blockade by either HPV or inhibitors released this control, compromised HRR and increased response to PARP inhibition. Antagonizing miR-182 rescued the homologous recombination deficit in HPV+ cells. Loss of TGFβ signaling unexpectedly increased error-prone, alternative end-joining repair.

**Conclusions**: HPV-positive HNSCC cells are unresponsive to TGFβ. Abrogated TGFβ signaling compromises homologous recombination and shifts reliance on alt-EJ repair that provides a mechanistic basis for sensitivity to PARP inhibitors. The effect of HPV in HNSCC provides critical validation of TGFβ’s role in DNA repair proficiency and further raises the translational potential of TGFβ inhibitors in cancer therapy.

## Introduction

The prognosis for patients with head and neck squamous cell carcinomas (HNSCC) is associated with anatomical location, tumor characteristics and patient history (1). Oropharyngeal squamous cell carcinoma patients whose cancer is positive for human papillomavirus (HPV+) have a markedly better prognosis than patients with compared to those whose tumors are HPV-negative (HPV-) (2). Recognition of this differential survival as a function of HPV in HNSCC has motivated dual goals to identify the mechanism by which HPV alters the DNA damage response (DDR) and to search for a means to achieve similar outcomes in HPV-HNSCC. HPV+ cell lines are more sensitive to cytotoxic agents (3,4) and exhibit decreased DNA repair capacity that may explain increased response to chemoradiation, of which ionizing radiation and cisplatin are standard-of-care in HNSCC (5,6). Identification of a specific DDR deficiency represents a vulnerability that can be exploited by cytotoxic therapy (7).

The cell cycle machinery is necessary for HPV replication (8). Transforming growth factor (TGFβ) profoundly suppresses epithelial cell proliferation. Studies in cervical cancer shows that HPV encoded E5, E6 and E7 proteins bind to TGFβ receptors and signal transducers, resulting in their degradation, thereby releasing infected cells to proliferate (9). Although TGFβ is considered a canonical tumor suppressor, most cancers both produce abundant TGFβ and maintain competent TGFβ signaling (10). TGFβ binds a ubiquitous heteromeric complex of type I (TβRI) and type II receptors whose serine/threonine kinase activity initiate canonical signaling cascade via phosphorylation of SMAD (signaling mother against decapentaplegic peptide) 2 and 3. SMAD 2/3 phosphorylation enables formation of SMAD4 heteromeric complexes that bind SMAD-binding gene elements to control transcription. Notably, SMAD4 is frequently mutated or deleted in HPV-HNSCC (11), but whether HPV affects TGFβ signaling in HNSCC has not been examined.

Of its myriad roles, TGFβ multiple action to maintain genomic stability is among the most poorly appreciated. Yuspa and colleagues first reported in 1996 that *Tgfb1* null keratinocytes exhibit profound genomic instability (12). Genetic deletion of *Tgfb1* or inhibiting TGFβ activity or signaling compromises DNA damage recognition and execution of canonical cell fate decisions, i.e. repair, cycle arrest and cell kill in vivo or in vitro (13). This is in part because TGFβ signaling is necessary for ataxia telangiectasia mutated (ATM) kinase activity during DNA damage (14) and ligase IV (15); both are important components of classical non-homologous end joining (NHEJ) repair of DNA damage. TGFβ also regulates breast cancer early onset 1 (BRCA1) (16,17), which is a major component of homologous recombination repair (HRR). Together, the multiple mechanisms by which TGFβ controls DNA damage components raise the potential translational utility of pharmaceutical blockade of TGFβ signaling during cancer therapy (18,19). Consistent with this, TGFβ inhibition promotes response to radiation therapy and/or chemotherapy in preclinical models of breast (20), brain (21–24) and lung (25) cancer.

Here, we investigated whether HPV affects TGFβ signaling in HNSCC and if so, whether loss of TGFβ signaling underlies the responsiveness of HPV+ HNSCC to cytotoxic therapy.

## Materials and Methods

***Cell lines*** HNSCC cell lines (see Supplemental Figure 1B) were cultured in Dulbecco’s modified Eagle’s medium (DMEM), supplemented with 10% fetal bovine serum (HyClone), GlutaMax (Thermo Fisher), HEPES and 100 IU/ml Streptomycin-Penicillin (all Sigma-Aldrich unless otherwise indicated). Detroit562, SCC090, SCC154 and FADU were purchased from the American Type Culture Collection (Rockville, MD). SCC47 and SCC104 were purchased from Millipore Sigma. HPV status of these cell lines were validated by PCR(26). All cells were maintained in a humidified incubator at 37°C and 5% CO_2_. All cell lines were tested mycoplasma free and validated for their DNAss identities by microsatellite markers for authentication (IDEXX). Cells were used within 10 passages with restricted culture time (max. 6 weeks) for all experiments after defrosting. Cells were maintained in exponential growth phase before sample preparation for each experiment.

***Patient Derived Xenografts*** Balb/c nude mice were purchased from The Jackson Laboratory. All mouse experiments were approved by and performed according to the guidelines of the Institutional Animal Care Committee of the Vall d’Hebron Research Institute in agreement with the European Union and national directives. Subcutaneous inoculation of 1×10^6^ cells SAS in the flank was performed in 7 week old mice with 8 mice per group for treatments. Treatments began at day 10 post inoculation after randomization of tumor sizes. LY2157299 (15 mg/Kg) was given by oral gavage twice a day and olaparib (5 mg/Kg) was given by oral gavage once a day. Tumor volume was measured daily using caliper method and weight monitored every day.

***Treatments*** TGFβ1 (500 pg/ml; R&D Systems, Inc.) was given in serum-free medium. Small molecule inhibitors of the TβRI kinase, LY364947 (Calbiochem) and LY2157299 (Galunisertib; SelleckChem) were used at 2 ⁊M; a pan-isoform TGFβ neutralizing monoclonal human antibody, GC-1008 (Fresolimumab), was used at 10 ⁊g/ml. All TGF⁊ inhibitors were given for 24-48 hr prior to other treatments. PARP1/2 inhibitor olaparib (LC Laboratories) and PARP1 inhibitor AG14361 (SelleckChem) were used at 10 ⁊M and 1 ⁊⁊M, respectively. ATM inhibitor KU55933, ATR inhibitor AZD6738, and DNA-PK inhibitor KU57788 (all SelleckChem) were used at 5 ⁊M, 0.5 ⁊M and 1 ⁊M, respectively. Inhibitors against PARP, ATM, ATR, and DNA-PK were given 1 hr prior to irradiation. All the above compounds were dissolved in dimethyl sulfoxide (Sigma-Aldrich) and stored at −20°C for up to 6 months with protection from light. They were aliquoted for maximum 3x use to avoid thaw-freeze cycles. Cisplatin (Santa Cruz) was dissolved in 0.9% sodium chloride in water to achieve 4 mM stock solution, stored at 4°C for up to 1 month with protection from light. Cells were irradiated to the indicated doses using 250 kV X-ray.

***Cell survival and proliferation assays*** Cells grown to 70 % confluence with 24 hr pre-treatment of LY364947 before irradiation. Cells were trypsinized 3 hr after irradiation and diluted into the appropriate densities with single cell suspensions, which were seeded in triplicates at 2 dilutions into 6-well cell culture plates. For each experiment, 3 or more independent biological repeats were performed. Cell seeding numbers were selected based on prior tests of each cell line for plating efficiency and radiosensitivity to yield colony numbers in the range from 20 to 100 colonies per well. Colonies were allowed to grow for a cell line dependent time (10 - 20 days) followed by fixing with acetic acid/ methanol solution (volume ratio, 1:3) and staining with 0.5% crystal violet. Colonies containing at least 50 cells were scored under a bright field microscope.

Plating efficiencies were calculated as colonies per number of cells plated, and surviving fractions as ratios of plating efficiencies for irradiated and unirradiated cells. For each radiation dose, the number of colonies obtained from 3 wells was averaged. These mean values were corrected according to plating efficiency and used to calculate the cell survival for each dose level. Linear-quadratic formula (LQ): ln(SF) = −(αD+βD^2^), was used to fit survival curves and calculate dose enhancement ratio at 10% surviving fraction (DER10%, i.e. DER).

The number of cells plated under each condition was selected in cell proliferation assay so that all cultures reached similar degrees of confluence (<80%) at experiment termination. Relative response was calculated as number of treated cells divided by untreated control 5-7 days (consistent time were used for repeated experiments in each assay) after treatments measured by SYTO60 assay, a fluorescent nucleic acid stain, as described (27), an ATP based luminescence assay CellTiter-Glo (Promega, Madison, WI, USA) following the manufacturer’s protocol, or cell number counting with automated counter (Countess, ThermoFisher). SRF_2Gy_ was calculated as the ratio of the cell fraction for IR alone and the cell fraction for combined drug/IR effect, corrected for the cell fraction for drug alone (27).

***Flow cytometry assays*** Cell cycle distributions were investigated by flow cytometry analyses of DNA staining with propidium iodide (PI; Sigma-Aldrich) according to standard protocol. Cells were collected and washed twice with PBS, and then fixed in 80% cold ethanol. Cells were kept in −20°C until analyses. Ethanol fixative was removed after spin-down cells. After washing twice in PBS, cells were incubated with 1 mg/ml ribonuclease (RNase) and 500 ⁊g/ml PI, and then analyzed by flow cytometer (Calibur, BD Biosciences). Apoptotic cells were determined by annexin V flow as previously described (27). Cells were harvested two days after treatment. Floating cells in the medium were also collected. Cells were spun-down, washed twice with cold PBS, resuspended in annexin binding buffer with cell density adjusted to 10^6 /ml, and pushed through a mesh capped tube to further separate cells to single cell suspension. Cells were stained with annexin V with Alexa Fluor 488 conjugate and PI according to the manufacturer’s protocol (ThermoFisher). Data were analyzed by FlowJo.

***Tissue microarray*** The tissue microarray (TMA) was generated within a retrospective study of the German Cancer Consortium Radiation Oncology Group (DKTK-ROG) and consists of up to three tissue cores of the untreated tumor and one or two cores of the adjacent tissue from 221 patients who were diagnosed with locally advanced HNSCC. All patients received postoperative, state-of-the-art cisplatin-based radiochemotherapy, and their inclusion criteria have been described previously (2). For the present study, tissues of 194 patients were evaluable and their characteristics are summarized in Supplemental Figure 1G. HPV DNA analyses of the tumors were part of a previous publication, methods were described previously (2).

***Explant specimens*** HNSCC PDX were collected from mice in which tumors were grown. Primary patient tumor tissues were collected during surgery. All specimens were kept in DMEM and transported on ice with HPV status blinded to experimenters. Explants were established within 5 hr on floating rafts as described (28). On day 1 explants were dissected and distributed for treatments with or without T⁊RI for 24 hr prior to irradiation with the indicated dose. At selected time points, samples were embedded in optimum cutting temperature compound (Sigma Aldrich) and frozen on dry ice. Samples as well as slides after sectioning were kept at −80°C. Patient characteristics for primary tissues are shown in Supplemental Figure 6H.

***Immunofluorescence and microscopy*** Staining and visualization of DNA damage repair foci was performed as previously described (25). Cells were plated on chamber slides with sub-confluent cell density until fixing with 4% paraformaldehyde (PFA). Cells were then permeabilized with 0.5% TritonX-100, and blocked with 0.5% Casein in PBS. Cells were incubated with γH2AX mouse monoclonal antibody (EMD Millipore) at 1:500 dilution, RAD51 rabbit polyclonal antibody (Santa Cruz) at 1:200, Geminin rabbit polyclonal antibody (Abcam) at 1:200, or phospho-SMAD2 on serine 465/467 at 1:200 (Cell Signaling) at room temperature for 2 hr or 4°C overnight. For HNSCC PDX and primary patient tumor tissue analyses, slides were defrosted at room temperature and fixed with 4% PFA for 15 minutes. Nonspecific sites were blocked in 0.5% casein in PBS for 1 hr at room temperature. Primary antibodies against γH2AX, RAD51, 53BP1 (Bethyl), Geminin, or p-SMAD2 were incubated overnight at 4°C in a humidified chamber. After 3x washes with PBS, secondary antibody including donkey anti-rabbit IgG (Alexa Fluor 488/555, Invitrogen) and donkey anti-mouse IgG (Alexa Fluor 488/555, Invitrogen) was incubated for 1 hour. Cell nuclei were counterstained with 4′,6-diamidino-2-phenylindole, dihydrochloride (DAPI). Slides were mounted in Vectashield (Sigma). A 40X objective with 0.95 numerical aperture was used on a Zeiss Axiovert equipped with epifluorescence. In-home developed macros or Findfoci plugins (29) in the open source platform Fiji-ImageJ (NIH, Bethesda, MA) were used for image analyses of 8-bit images for each channel of fluorescence. The DAPI channel was used to generate region of interest in 5 or more images randomly taken based on nuclear dye alone. At least 100 cells were analyzed for each treatment. For analysis of radiation-induced foci, spontaneous foci from sham-treated controls were subtracted unless otherwise noted.

For the patient tissue array (30), intensity of p-SMAD2 was scored by three observers blinded to tissue identity according to the following criteria: level 1, no p-SMAD2 signal (negative −); level 2, a little p-SMAD2 signal (positive +); level 3, a medium p-SMAD2 signal (positive ++); level 4, a strong p-SMAD2 signal (positive +++), as illustrated in Supplemental Figure 1F. The independent scores were averaged.

***Western blotting*** Proteins from exponentially growing cells were extracted on ice by radioimmunoprecipitation assay buffer with protease and phosphatase inhibitor cocktails (Sigma Aldrich), quantified, and 40-100 μg was electrophoresed on 4-15% gradient gels from BioRad and transblotted on fluorescence-optimized PVDF membrane (Merck Millipore). The membrane was blocked in blocking buffer and probed with one of the primary antibodies including BRCA1 with antibodies against the N-terminus (a gift from Dr. Chodosh) or C-terminus (Santa Cruz), FOXO3 (Cell Signaling), ATM (GeneTex), p-ATM (Ser-1981, Abcam), or β-actin (Abcam). The membrane was washed 3x 5 min with 0.05% Tween-20 in TBS (TBST), followed by incubation with secondary antibodies (Odyssey) in dark for 1 hr at room temperature. The membrane was washed 3 times with TBST again and scanned on the Odyssey LICOR system.

***Immunoprecipitation*** Cell lysates were prepared using specialized lysis buffer (Thermo Fisher) and protein G Sepharose beads were used for immunoprecipitation following the manufacturer’s instructions (GE Healthcare) using rabbit monoclonal antibody to FOXO3 (Cell Signaling) or ATM (GeneTex). The presence of ATM and p-ATM, or FOXO3 was probed by immunoblotting.

***qRT-PCR*** Total RNA was extracted from samples using the TRIzol reagent (Invitrogen) and miRNAeasy Mini Kit (Qiagen) according to manufacturer’s instructions. cDNA was synthesized from 1 μg of RNA using the SuperScript III First-Strand Synthesis System (Invitrogen). Real-time quantitative PCR was performed using SYBR Green Mix (Applied Biosystems), according to the manufacturer’s instructions. Primers with published sequences (17) were purchased from Sigma-Aldrich. Expression of genes of interest was normalized against expression of *GAPDH*. qRT-PCR analyses of miR-182 was performed using miRNA-specific TaqMan MicroRNA Assay Kit; 12.5 ng of total RNA was reverse transcribed using the corresponding RT Primer and the TaqMan MicroRNA Reverse Transcription Kit (Applied Biosystems). TaqMan PCR primers and the TaqMan Universal PCR Master Mix were added to reverse transcribed products for PCR analyses (Applied Biosystems). RNU44 was used for normalization of input RNA/cDNA levels. Data were analyzed using the comparative Ct method for quantification of transcripts.

***Transfections*** Exponentially growing cells were transfected with 50 nM *FOXO3* siRNA (sense: CUCACUUC GGACUCACUUAtt; antisense: UAAGUGAGUCCGAAGUGAGca; Silencer^®^ Select Pre-designed (ThermoFisher) or scrambled non-targeting siRNA. 50 nM of miR-182 mimic or 200 nM of miR-182 inhibitor (i.e. anti-miR) were transfected into cells overnight in parallel with non-targeting miRNA control (mirVana, Thermofisher). Published *FOXO3*-expressing plasmid with pCMV5 vector (Addgene) was transfected overnight into SAS cell line (31). Validated shRNA Lentiviral transduction particles with pLKO.1 vector (Sigma Aldrich) were used to knock-down SMAD4 or POLQ in SAS cell line, with control cells prepared in parallel using vector control lentiviral particles. Cells were infected with 8 ⁊g/ml polybrene overnight and grown in 2 ug/ml puromycin for selection. Lipofectamine 3000 Transfection Kit (Invitrogen) was used according to the manufacturer’s instructions. Western blotting to determine protein levels of targeted genes and subsequent experiments were carried out 48 hours after transfection.

***DSB repair reporter assay*** DNA repair reporters of HRR (pDRGFP) or alt-EJ (EJ2GFP-puro) were stably integrated into clones of the SAS cell line using the published plasmid constructs (32,33). The expression of the integrated GFP gene expression is selectively activated in these clones was then used to determine the frequency of events from HRR- or alt-EJ, following the generation of DSB in the reporter constructs by I-SceI endonuclease. Agarose electrophoresis analyses using a variety of restriction enzyme digestions including I-SceI was used to verify the identities of these plasmids (data not shown). To establish reporter cell clones, linearized plasmids with appropriate restriction enzyme were transfected into exponentially growing SAS cells using Lipofectamine 3000 in Opti-MEM. Cells were then selected in 2 ug/ml puromycin containing medium. Single cell clones were established in 96-well plates with one cell per well. pQCXIH-I-SceI was packaged into retroviral vectors for infection as previously described (34). Infected cells were selected in culture medium with 400 ⁊g/ml hygromycin for 10 days. Medium, as well as treated drugs, were replaced every 2-3 days. Successful I-SceI infection showing GFP fluorescence was observed under fluorescent microscope. For analyses, the treated cells were trypsinized and maintained on ice before flow cytometry used to determine the fraction of GFP-positive cells. Propidium iodide staining was used to exclude dead cells from analyses.

***Chromosome aberration analysis*** Chromosome aberrations were analyzed using the standard metaphase spread assay. Cells at exponential growth were plated one day before treatments. At 48 hr post radiation, colcemid (Sigma) were added for 2 hours to impede mitosis; mitotic cells were then collected, washed, and suspended in warm hypotonic (75 mM KCl + 10% FBS) solution for 20 min at 37 °C. Cells were fixed in 3:1 methanol: glacial acetic acid at 4°C overnight. Cell suspensions were dropped onto warm slides at 50 °C, and air dried for 24 hours afterwards. Chromosomes were stained with DAPI in Vectashield (Sigma). Metaphases were imaged using Zeiss AxioImager 2 at 63x magnification. Chromosome aberrations were scored manually by three blinded experimenters from at least 100 metaphases in each treatment conditions of three independent experiments.

***Comet assay*** Comet assay was performed according to the manufacturer’s protocol (Trivigen). Tumor samples after treatments were dissected into small pieces for enzymatic disaggregation including 1.5 mg/mL collagenase and 0.25% trypsin at 37°C for 1hr (Thermo Fisher). Cell strainer with 40 μm micron pores was used to obtain single cell suspension for comet assay. Single-cell gel electrophoresis was performed at 19 V, 40 min. SYBR Gold-stained DNA comets were imaged with a Zeiss fluorescence microscope. Images were analyzed using Fiji-ImageJ with OpenComet (35). The Olive tail moments of minimum 100 cells were determined in each sample by OpenComet.

***TCGA and gene expression analyses*** Clinical and gene expression data of HNSCC was obtained from cBioPortal for Cancer Genomics (http://www.cbioportal.org/public-portal/) and from the corresponding TCGA publication (36). A chronic TGFβ signature was defined using a non-malignant epithelial cell line, MCF10A, that were exposed to TGFβ or LY364957 for 7 days and analyzed using Affymetrix gene expression microarray to identify genes with more than 2-fold change that were reciprocally regulated following chronic TGFβ treatment versus signaling inhibition. The RNAseq TCGA data was used for correlation (based on Pearson’s correlation coefficient), hierarchical clustering (based on Euclidean distance) and survival analyses. The association with survival was computed in R software (*survfit* package) using multivariate Cox regression analyses adjusted by age, gender, tumor stage and smoking status. The ssGSEA scores were computed using the GSVA package in Bioconductor (37).

***Statistical Analyses*** The data were analyzed by GraphPad Prism 6. Bars or data points represent means based on at least 3 independent experiments with error bars indicating standard error or medians with 95% confidence intervals as indicated in figure legends. Two-tailed Student’s t-test or Mann-Whitney U test was used for statistical comparisons and considered as significantly different at *P* < 0.05. F-test was used to test the clonogenic survival effect after treatments by assessing coefficients α and β in derived formula by Linear-Quadratic fitting. P-values are indicated in the figures as follows: ***, *P* < 0.005; **, *P* < 0.01; *, *P* < 0.05.

***Study approval*** Patient specimens were obtained with written consent from each patient in accordance with the ethics guidelines for research in the US (protocols # 14-15342, approved by the IRB committee of UCSF) and in Germany (protocol# EK299092012, approved by the IRB committee of the DKTK partner site and subsequently by the IRB committees of all other DKTK partner sites).

## Results

### TGFβ signaling is decreased in HPV+ HNSCC

We first sought to determine using data from The Cancer Genome Atlas (TCGA), of which 13% (n=36) are HPV+ (36) whether TGFβ signaling is associated with HPV status in HNSCC. We derived a chronic TGFβ expression signature in non-malignant MCF10A epithelial cells treated for 7 days with either TGFβ or a small molecule inhibitor of TGFβ type I receptor (TβRI) kinase, LY364947 (Supplemental Figure 1A). Thirty genes that were induced by at least 2-fold by TGFβ and blocked by LY364957 were expressed in HNSCC and used for supervised clustering. Almost all HPV+ cancers were clustered in the clade with low expression of TGFβ induced genes (Figure 1A). For example, expression of canonical TGFβ genes CTGF, FN1, POSTN and SERPINE1 2, were low in the right most dendrogram arm in which HPV+ HNSCC were clustered compared to the high expression in leftmost arm that is nearly exclusively HPV− HNSCC. The negative correlation suggests that TGFβ signaling is decreased in HPV+ HNSCC.

**Figure 1.**
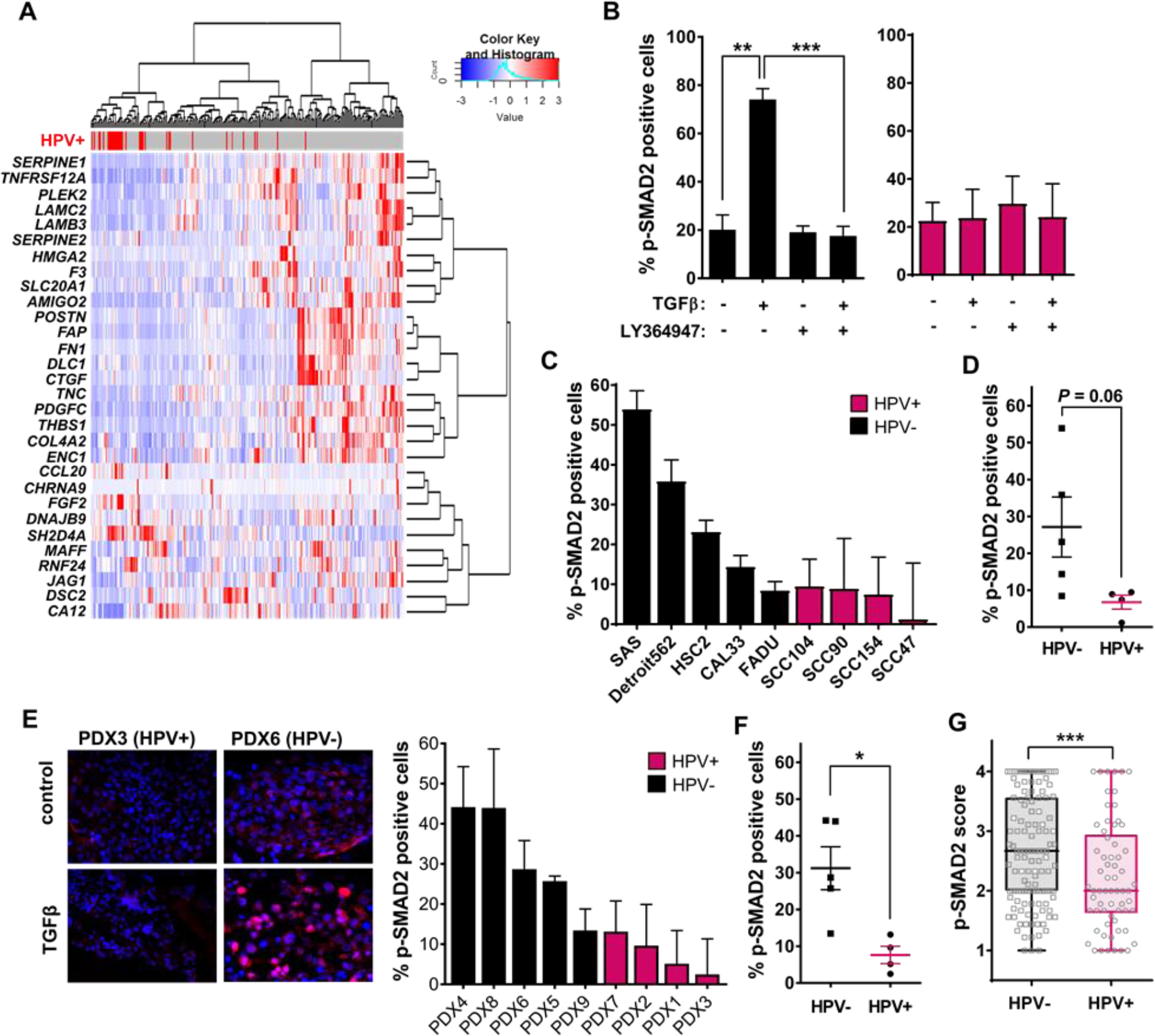
TGFβ signaling is defective in HPV+ HNSCC. **(A)** TCGA tumor sample clustering of 243 HPV− and 36 HPV+ HNSCC using a gene-set signature upregulated by TGFβ signaling. **(B)** Percentage of p-SMAD2 positive cells after treatment with TGFβ or TβRI inhibitor LY364947 for a representative HPV− cell line, SAS (black), and a HPV+ cell line, SCC47 (red). Two-tailed Student’s t-test; **, *P* < 0.01; **, *P* < 0.005. **(C)** The percentage of p-SMAD2 positive cells following TGFβ treatment of HPV− cell lines (black) and HPV+ cell lines (red). **(D)** Data from panel C grouped according to their HPV status. **(E)** Representative images of p-SMAD2 immunostaining of HNSCC PDX (left panel). The percentage of p-SMAD2 positive cells induced by TGFβ in HPV− (black) and HPV+ (red) PDX (right panel). **(F)** Data from panel E grouped according to HPV status. **(G)** Distribution of p-SMAD2 scores as described in methods for 130 HPV− and 65 HPV+ HNSCC. Panel D, F and G, Mann-Whitney U test; *, *P* < 0.05; ***, *P* < 0.005.

Next, we sought to validate the association between HPV status and TGFβ signaling in a panel of HNSCC cell lines (Supplemental Figure 1B). TGFβ type I receptor (TβRI) mediated phosphorylation of SMAD 2 and 3 initiates TGFβ signal transduction. Protein levels of SMAD 2 and 3 and TGFβ type I and II receptors were similar among 3 HPV-and 2 HPV+ cell lines (Supplemental Figure 1C), however none of the cell lines were growth inhibited by TGFβ treatment. We assessed canonical signaling by localizing nuclear phosphorylated SMAD2 (p-SMAD2) in cells treated briefly with exogenous TGFβ. TGFβ treatment increased p-SMAD2, which was blocked by treating cells with TβRI kinase inhibitor LY364957 in an HPV− cell line, as expected. In contrast, TGFβ treatment did not increase in p-SMAD2 in an HPV+ cell line (Figure 1B). As a group, HPV+ cell lines showed minimal response to TGFβ compared to HPV− cell lines (Figure 1, C and D).

We then tested p-SMAD2 response following acute TGFβ treatment of explants of HNSCC patient-derived xenografts (PDX) (Supplemental Figure 1D). As found in the cell lines, TGFβ elicited more robust nuclear p-SMAD2 in HPV− PDX compared to HPV+ PDX (Figure 1E). Overall, the levels of p-SMAD2 of HPV+ PDX (n=5) were significantly less than that of HPV− PDX (n=4) (Figure 1F). Lastly, we performed semi-quantitative evaluation of p-SMAD2 in a human HNSCC tissue microarray consisting of 194 cases (Supplemental Figure 1E) (30). HPV+ cancers exhibited significantly lower p-SMAD2 levels than did HPV− cancers (*P* = 0.002; Figure 1G). Levels of p-SMAD2 in normal epithelium were not affected by the HPV status of the adjacent cancer (Supplemental Figure 1F). Together these analyses of TCGA, cell lines, PDX explants and tumor specimens provide comprehensive and compelling evidence that TGFβ signaling is significantly compromised in human HPV+ HNSCC.

### Impaired TGFβ signaling associates with better response to radiation and chemotherapy

Our prior work showed that inhibiting TGFβ signaling with LY364947 increased the radiosensitivity of breast, brain and lung cancer cell lines (19,20,22). A short term assay that reflects both cell cycle delay and cell kill by measuring cell number 5 days after exposure to 2 Gy (27) showed that HPV+ HNSCC cell lines appeared more sensitive to radiation than HPV− HNSCC cell lines (Figure 2A), and was significant different as a function of HPV status (*P* = 0.02; Figure 2B). We then determined radiosensitivity using the response ratio at 2 Gy (SRF_2Gy_), as previously reported (27). Inhibition of TGFβ signaling using LY364947 radiosensitized all HPV-cell lines (i.e. SRF_2Gy_ >1), whereas only 1 (SCC154) of 4 HPV+ cell lines was radiosensitized (Figure 2C). As a group, LY364947 treatment significantly increased radiosensitivity of HPV-cell lines compared to HPV+ HNSCC cell lines (*P* < 0.05; Figure 2D). Notably, the degree of TGFβ responsiveness measured by p-SMAD2 and radiosensitivity measured by SRF_2Gy_ were significantly correlated (*P* = 0.04; Supplemental Figure 2A).

**Figure 2.**
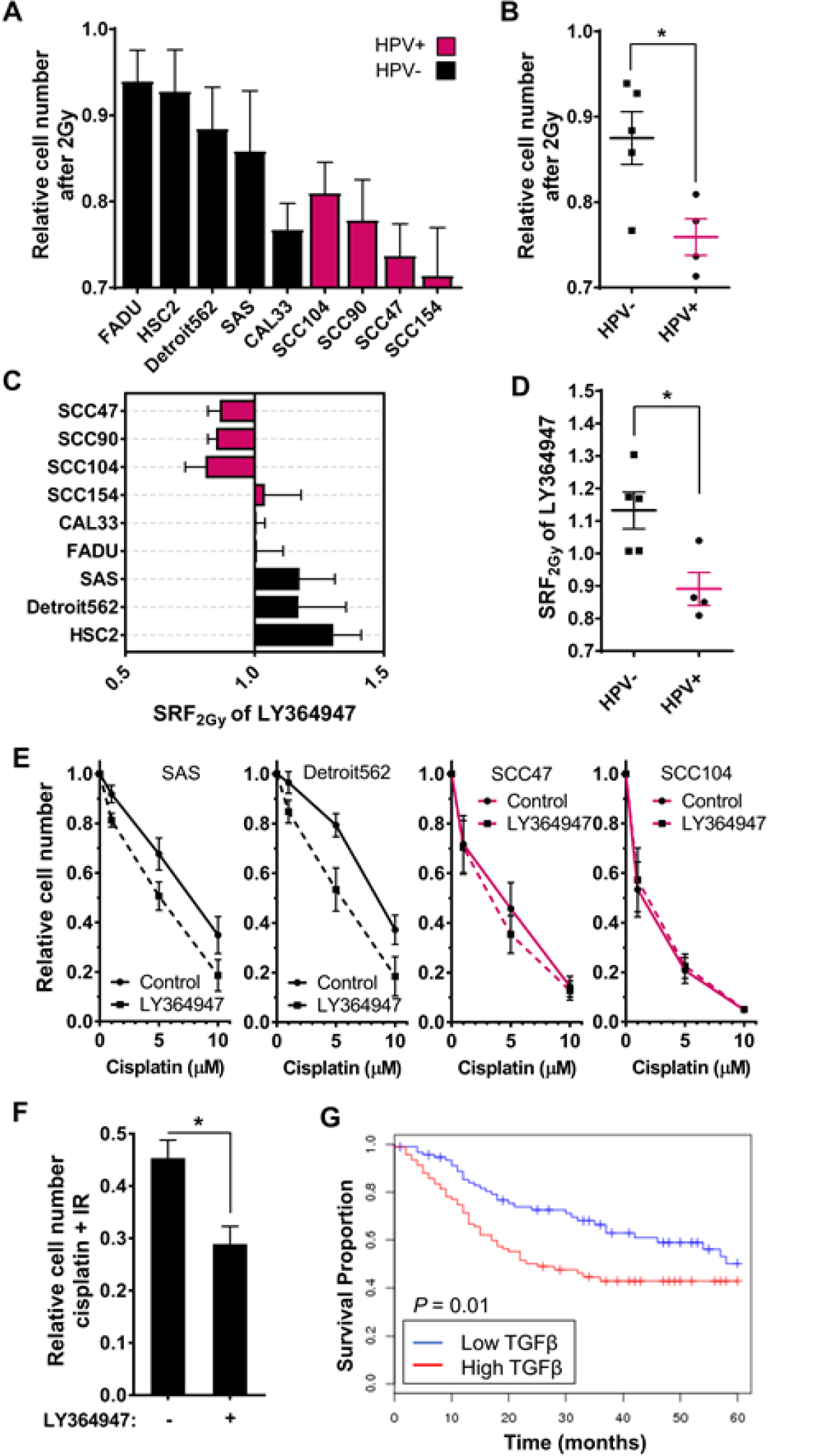
HNSCC sensitivity to IR or cisplatin correlates with TGFβ signaling status. **(A)** Relative cell number after irradiation (2 Gy) measured by a 5 day cell viability assay. HPV− (black), HPV+ (red). **(B)** Relative cell number after irradiation (2 Gy) from panel A shown as HPV− versus HPV+ cell lines. Two-tailed Mann-Whitney U test; *, *P* < 0.05. **(C)** SRF_2Gy_ of TβRI inhibitor, LY364947, was measured on the HNSCC cell lines. **(D)** SRF_2Gy_ values from HPV− cell lines versus HPV+ lines. Two-tailed Mann-Whitney U test; *, *P* < 0.05. **(E)** Relative cell number after cisplatin treatments compared to sham-treated control were measured in a 5 day cell proliferation assay. **(F)** Relative cell number after combination treatments with cisplatin (5 ⁊M) for 1 hr and 2 Gy, with or without LY364947. Two-tailed Student’s t-test; *, *P* < 0.05. **(G)** Kaplan-Meier survival analyses of HNSCC patients from TCGA with top (i.e. high TGFβ; red) or bottom one-third (i.e. low TGFβ; blue) according to expression levels of TGFβ-upregulated genes in Supplemental Figure 1A. Log-rank test, *P* = 0.01.

We confirmed radiation sensitivity by classic clonogenic assay. LY364947 also significantly increased radiosensitivity of the HPV− cell line SAS, but did not affect the radiosensitivity of HPV+ cell line SCC47. Note that SCC47 displayed higher intrinsic radiosensitivity compared to SAS (Supplemental Figure 2B). The response to TGFβ inhibition by LY364947 was confirmed by employing two other drugs currently in cancer clinical trials to inhibit TGFβ signaling: a TβRI inhibitor, LY2157299 (Galunisertib) and TGFβ pan-neutralizing antibody, GC1008 (Fresolimumab). Both drugs radiosensitized HPV− SAS cells, (Supplemental Figure 2C), but neither inhibitor increased radiosensitivity of HPV+ cell line SCC47 (Supplemental Figure 2D). TGFβ inhibition by LY364947 also sensitized HPV-cancer cells to cisplatin, but not HPV+ cells, which were intrinsically more sensitive to cisplatin (Figure 2E). Moreover, HPV-cells exposed to cisplatin in combination with IR were further sensitized treatment with LY364947 (Figure 2F). The increased cytotoxic response of HNSCC cells in which TGFβ signaling is impaired should improve patient outcomes. Consistent this, survival of patients in the TCGA cohort whose cancers exhibit low TGFβ activity was significantly better (Figure 2G).

### MiR-182 mechanism by which loss of TGFβ signaling in HPV+ HNSCC impairs DDR

Our prior work showed that *Tgfb1* genetic deletion impedes ATM auto-phosphorylation and phosphorylation of its target histone H2AX (γH2AX) upon DNA damage (13,14). We hypothesized that defective TGFβ signaling in HPV+ HNSCC would phenocopy *Tgfb1*genetic deletion. We assessed frequency of IR-induced γH2AX foci as a function of LY364947, which did not change cell cycle distribution in either irradiated or control cells (Supplemental Figure 3A), nor did HPV status associate with significant difference in proliferation of PDX tissues (Supplemental Figure 3B). Pre-treatment with TβRI inhibitor greatly reduced γH2AX foci formation in irradiated HPV-cell lines but did not alter γH2AX foci frequency in HPV+ cell lines (Figure 3A). Note that the level of IR-induced γH2AX foci of HPV+ cell lines approximated that following TGFβ inhibition in HPV− cell lines. Similarly, fewer IR-induced γH2AX foci were evident in PDX from HPV+ HNSCC compared to HPV− HNSCC, in which LY364947 decreased the number of γH2AX foci (Figure 3B). The degree of TGFβ responsiveness measured by p-SMAD2 correlated with γH2AX foci per irradiated cell across HNSCC specimens and cell lines (Figure 3C).

**Figure 3.**
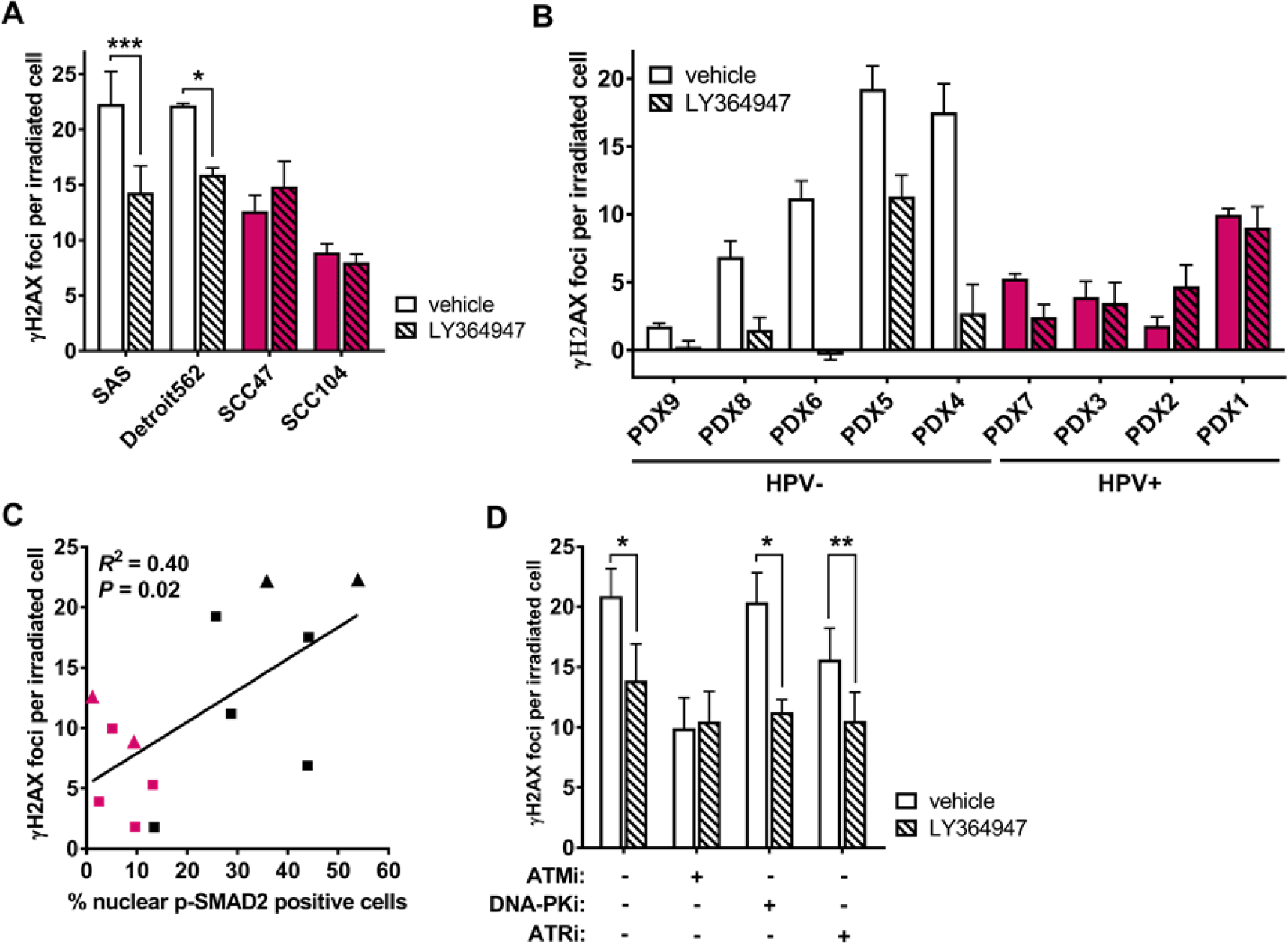
Impaired TGFβ signaling compromises DNA damage recognition through ATM. **(A)** Number of γH2AX foci per irradiated (2 Gy) HPV− cell lines (white) and HPV+ cell lines (red) at 30 min post irradiation pre-treated with (hatched) or without (open) TβRI inhibitor LY364947. **(B)** Number of γH2AX foci per cell in irradiated (2 Gy) HPV− (white) and HPV+ (red) PDX pre-treated with (hatched) or without (open) LY364947. **(C)** Correlation of percent nuclear p-SMAD2 positive cells following TGFβ treatment (from Figure 1, C and E) with γH2AX foci (from panel A and B) of HPV+ (red) and negative (black) cell lines (triangles) and PDX (squares). Linear regression analysis was used to calculate *P* value and *R*^*2*^. **(D)** Number of γH2AX foci per cell of irradiated (2 Gy) SAS cells treated with inhibitors against ATM (ATMi, KU55933), DNA-PK (DNA-PKi, KU57788), or ATR (ATRi, AZD6738) with or without LY364947. Two-tailed Student’s t-test; *, *P* < 0.05; **, *P* < 0.01; ***, *P* < 0.005.

Because histone H2AX can be phosphorylated by ATM, ataxia telangiectasia and Rad3 related (ATR), or DNA-dependent protein kinase (DNA-PK), we tested the contribution of these proteins using specific kinase inhibitors of each (Figure 3D). Pre-treatment with LY364947 further reduced IR-induced γH2AX foci on cells treated with DNA-PK or ATR inhibitor (KU57788 and AZD6738, respectively) but not in cells treated with ATM inhibitor (KU55933), consistent with our conclusion (14) that TGFβ signaling is necessary for ATM kinase activity (Supplemental Figure 3C).

The mechanism by which TGFβ affects ATM kinase activity has not been identified. Here, we investigated FOXO3 (forkhead box protein O3) because it is required for ATM auto-phosphorylation (38) and *FOXO3* is targeted by miR-182, which we have shown is regulated by TGFβ (17). *BRCA1* is also a target of miR-182 (17,39). Consistent with our prior studies, *BRCA1* expression is TGFβ responsive HPV− cell line but not an HPV+ cell line (Supplemental Figure 4, A and B). Consistent with this, TGFβ increased expression of *FOXO3* mRNA in a HPV− SAS cell line and TGFβ inhibition with LY364947 blocked this increase, while *FOXO3* expression was unaffected by TGFβ in the HPV+ cell line SCC47 (Figure 4A). We confirmed that both ATM and p-ATM were pulled down with FOXO3 immunoprecipitation in extracts from the HPV− SAS cell line (Figure 4B). To test whether FOXO3 was the missing link between TGFβ and ATM kinase, SAS cells were transfected with a *FOXO3* siRNA or scrambled control. In *FOXO3*-depleted cells, p-ATM did not change upon treatment with IR, LY364947, or their combination (Figure 4, C and D). We then overexpressed *FOXO3* in SAS cells (Figure 4E), which abrogated the inhibitor effect of LY364947 on IR-induced ⁊H2AX foci formation (Figure 4F). These data suggest that TGFβ regulation of FOXO3 is critical for its effect on ATM kinase activity.

**Figure 4.**
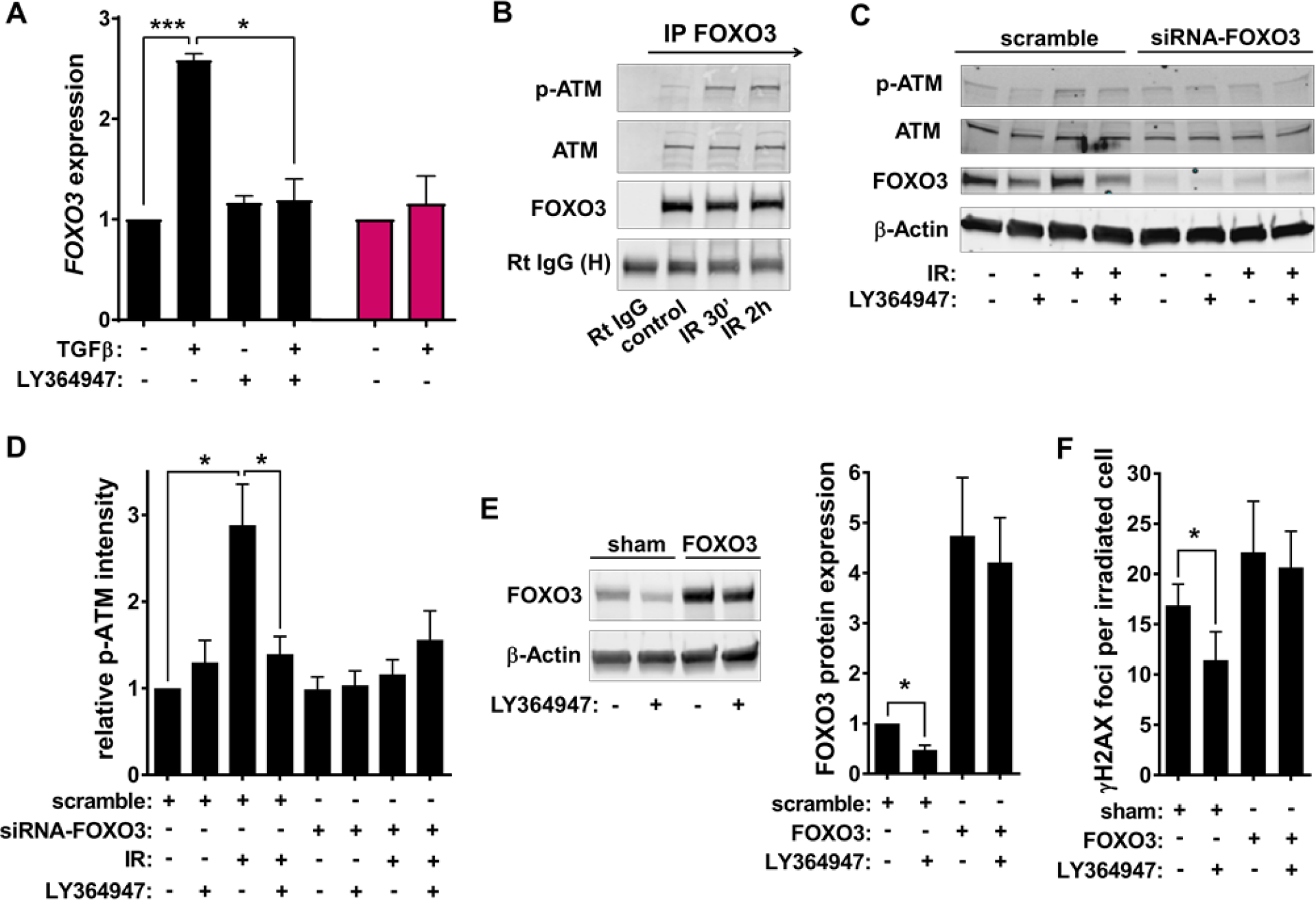
TGFβ regulation of *FOXO3* expression affects ATM kinase activity. **(A)** Levels of *FOXO3* by qRT PCR in HPV− SAS (black) and HPV+ SCC47 cell lines (red) treated with or without TGFβ, TβRI inhibitor LY364947, or combination of both. **(B)** Immunoprecipitation (IP) of FOXO3 in sham or irradiated (5 Gy) SAS cells and immunoblotted to detect for p-ATM and ATM. Control rabbit (Rt) IgG was used for sham-IP. Samples were also assayed for heavy chain of Rt IgG (H) as loading control. **(C)** Representative western blot images of p-ATM, ATM and FOXO3 from SAS cells transfected with scramble siRNA (i.e. scramble) or siRNA against *FOXO3* (i.e. siRNA-*FOXO3*) after irradiation (5 Gy) with or without LY364947. **(D)** Quantified protein expression of p-ATM normalized to total ATM and β-actin. **(E)** Protein bands of FOXO3 and β-actin measured on SAS cells, which were transfected with *FOXO3*-expressing plasmids or sham-transfected as control (sham), and then treated with or without LY364947. Protein expression was quantified from western blots. **(F)** Number of γH2AX foci per irradiated (2 Gy) cells at 30 min post irradiation pre-treated with or without LY364947. Cells at two days after transfection were used. Two-tailed Student’s t-test; *, *P* < 0.05; ***, *P* < 0.005.

Consistent with TGFβ regulation of miR-182, HPV+ cell lines exhibited higher levels of miR-182 compared to HPV− cell lines (Figure 5A). TGFβ suppressed miR-182 in a HPV-cell line but not a HPV+ cell line (Figure 5B). TGFβ-responsive, HPV− SAS cells were transfected with miR-182 mimic, anti-miR-182, or scrambled controls. Cell cycle distribution was not affected by modulating miR-182 expression (Supplemental Figure 4C) as published (39,40), and manipulation of miR-182 did not affect TGFβ induction of p-SMAD2 (Supplemental Figure 4D). MiR-182 mimic suppressed *FOXO3* and *BRCA1*, while anti-miR-182 increased both (Figure 5C and Supplemental Figure 4E). Cells transfected with anti-miR-182 and treated with LY364947 did not repress FOXO3 (Figure 5D) or BRCA1 (Supplemental Figure 4F). Since TGFβ control of miR-182 mediated FOXO3, we then examined whether functional DDR could be restored by inhibiting miR-182. Antagonizing miR-182 when TGFβ signaling was inhibited by LY364947 restored IR-induced phosphorylation of ATM (Figure 5E and Supplemental Figure 5A) and γH2AX foci formation (Figure 5F). Moreover, LY364947 radiosensitization of SAS cells was eliminated in cells expressing anti-miR-182 (Figure 5G and Supplemental Figure 5B).

**Figure 5.**
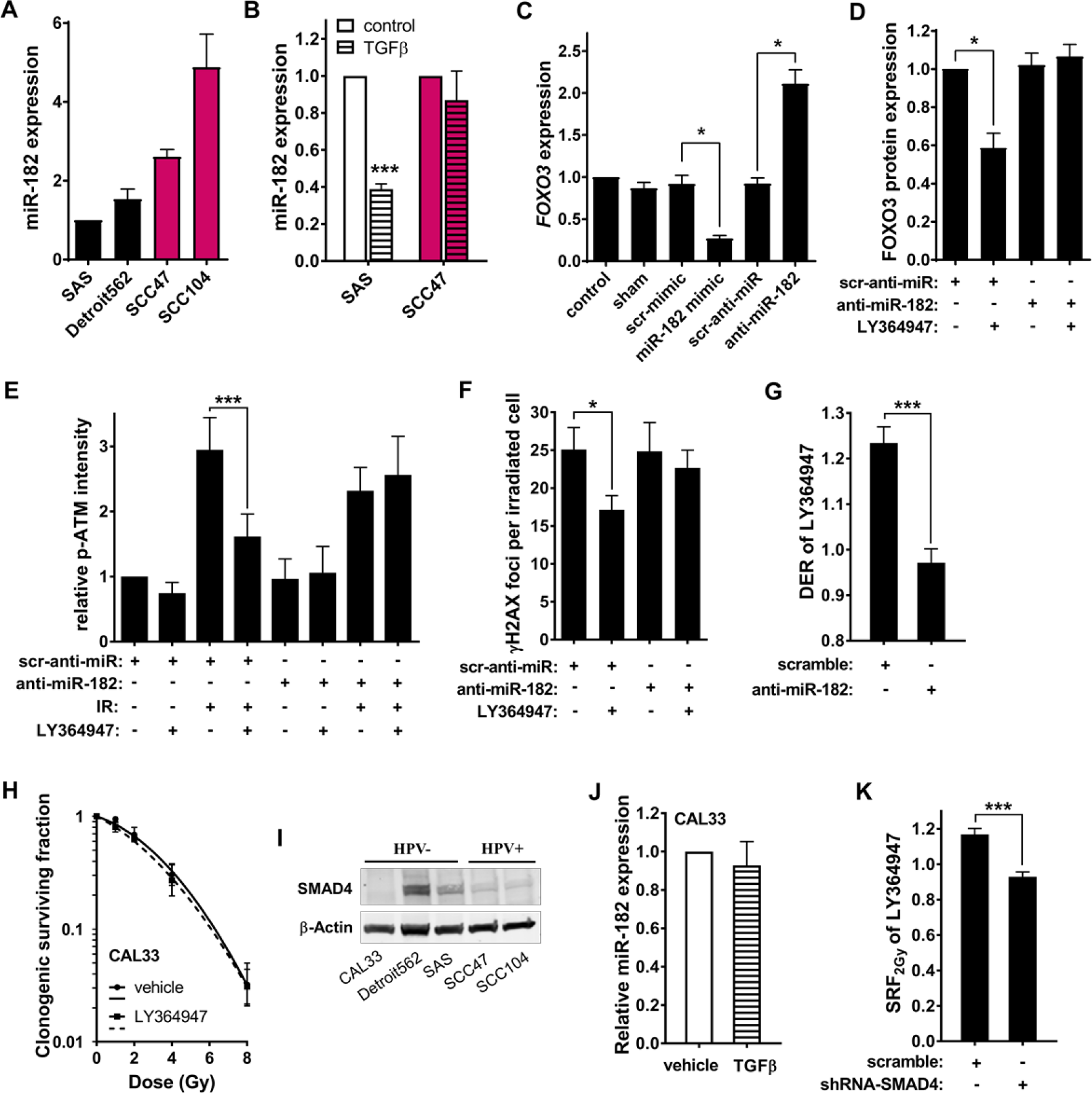
TGFβ controls DDR through miR-182 and SMAD4. **(A)** qRT-PCR analyses of miR-182 in HPV− (black) and HPV+ cell lines (red). **(B)** miR-182 expression in cells treated with (hatched) or without (open) TGFβ. **(C)** *FOXO3* mRNA levels in SAS cells transfected with miR-182 mimic, anti-miR-182, or scramble RNA for two days. **(D)** Quantified protein expression of FOXO3 with normalization to β-actin. SAS cells were transfected with scramble or anti-miR-182, and treated with or without TβRI inhibitor LY364947. **(E)** Quantified analyses of p-ATM protein bands, normalized to total ATM and β-actin, are shown as relative p-ATM intensities. SAS cells which were transfected with scramble RNA or anti-miR-182 and irradiated (5 Gy) with or without LY364947. **(F)** γH2AX foci per cell in irradiated (2 Gy) SAS cells transfected with scrambled or anti-miR-182 at 30 min post-irradiation. **(G)** Dose enhancement ratio (DER) of LY364947 measured by clonogenic survival assay on SAS cells transfected with scramble RNA or anti-miR-182. **(H)** Surviving fractions and fitted curves from clonogenic assay on CAL33 cells treated with or without TβRI inhibitor LY364947 prior to irradiation. (**I)** Protein bands of SMAD4 in five HNSCC cell lines. **(J)** qRT-PCR analyses of miR182 expression in CAL33 cells treated with or without TGFβ. **(K)** SRF_2Gy_ of LY364947 on SAS isogenic cells from lentiviral transduction with scramble or shRNA against SMAD4. Two-tailed Student’s t-test; *, *P* < 0.05; ***, *P* < 0.005.

Of the HPV− HNSCC cell lines, *SMAD4*-mutated CAL33 was not radiosensitized by LY364957 (Figure 5H). Consistent with this, SMAD4 protein was considerably reduced in Cal33 but was also reduced in HPV+ cell lines (Figure 5I). TGFβ did not suppress miR-182 in CAL33 (Figure 5J), but compared to TGFβ responsive SAS, mir-182 level was 2.4 fold less in CAL33, close to that of HPV+ cell lines. To test the hypothesis that TGFβ suppression of miR-182 requires SMAD4, we knocked down SMAD4 by shRNA in SAS cells (Supplemental Figure 5C). Doing so increased miR-182 expression (Supplemental Figure 5D) and eliminated the effect of LY364957 on the radiation response measured by SRF_2Gy_ (Figure 5K). We next examined the relationship between SMAD4 mutations and response to cisplatin using the data in the Sanger database (41). Consistent with loss of TGFβ signaling increasing cisplatin sensitivity in HPV+ cell lines (Figure 2E), cell lines in which SMAD4 was suppressed were significantly more sensitive to cisplatin (*P* < 0.005; Supplemental Figure 5E). Together these data led us to conclude that SMAD4 dependent suppression of miR-182 is the lynchpin by which TGFβ regulates DDR.

### Compromised TGFβ signaling impairs HRR proficiency

HRR repairs double strand breaks (DSB) in S and G2 phase when a homologous template is available, during which strand invasion is mediated by recombinase, RAD51, that binds to single-stranded DNA at the processed DSB (42). HRR is compromised in *SMAD4* mutant HNSSC (16). RAD51 foci formation is considered to be a specific marker for execution of HRR. To test whether loss of TGFβ signaling impeded HRR, we used immunostaining to detect RAD51 foci at its peak time (5-6 hr) in irradiated cell lines and PDX tumor explants. TβRI inhibition decreased RAD51 foci formation after IR in HPV− SAS cell line but had no effect in HPV+ SCC47 cell line (Figure 6A). Notably, antagonizing miR-182 in the HPV− SAS cell line also blocked the effect of TGFβ inhibition on RAD51 foci formation, consistent with HRR restoration (Supplemental Figure 6A). Most irradiated HPV− PDX explants (4/5) treated with TβRI inhibitor formed fewer RAD51 foci, however foci were barely above background in HPV+ PDX (Figure 6B). TβRI inhibition significantly reduced RAD51 foci formation in HPV− PDX explants and approached the low levels in HPV+ explants (Figure 6C). To validate TGFβ impact on HRR, we established HPV− SAS cells expressing constructs in which green fluorescent protein (GFP) is expressed if HRR occurs following an endonuclease-generated DSB (43). The frequency of HRR was decreased by approximately 50% when TβRI kinase was inhibited by either of two small molecules (Figure 6D), which was comparable to that observed following ATM inhibition by KU55933. These data indicate that HRR is compromised by inhibition of TGFβ signaling and is defective in HPV+ HNSCC.

**Figure 6.**
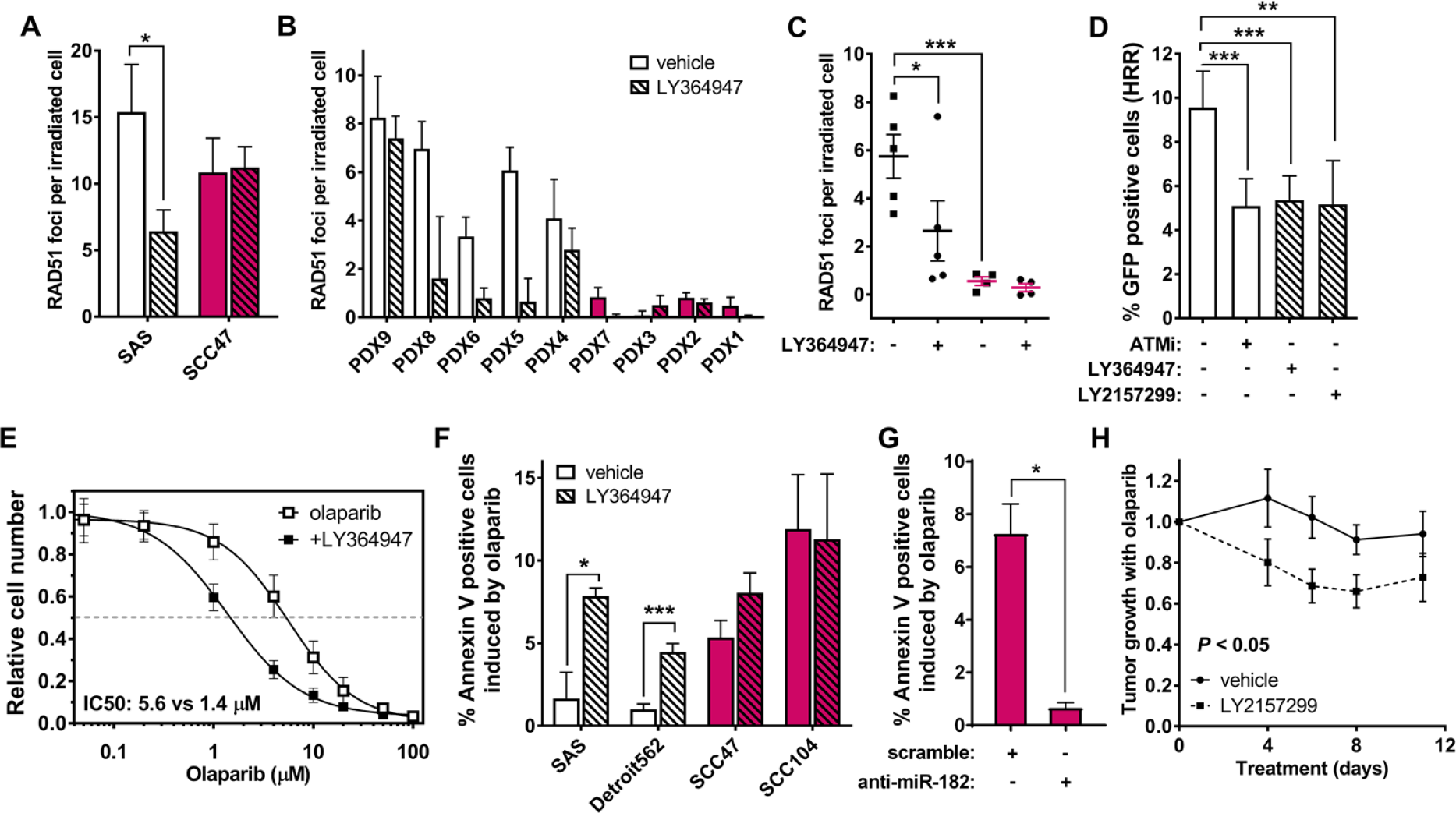
Decreased TGFβ signaling impedes DNA repair through HRR. **(A)** Number of RAD51 foci per cell at 6 hr post irradiation (4 Gy) in HPV− (white) SAS cell line and HPV+ (red) SCC47 cell line pretreated with (hatched) or without (open) LY364947. **(B)** RAD51 foci per cell of irradiated (4 Gy) PDX samples. **(C)** Data from panel B were grouped according to HPV status and LY364947 treatments. **(D)** Flow cytometry analyses of I-SceI-HRR reporter SAS cell clone treated with TβRI inhibitors, LY364947 and LY2157299, or ATM inhibitor (ATMi), KU55933. Percentage of GFP-positive cells after indicated treatments on I-SceI-HRR reporter cells. **(E)** Relative cell number after olaparib treatments with or without LY364947. Data were fitted with 4th polynomial sigmoidal curve for calculations of half maximal inhibitory (IC50) concentrations, which was 5.6 μM for olaparib alone and 1.4 μM for olaparib in combination with LY364947, respectively. **(F)** Percentage of annexin V positive cells induced by olaparib with (hatched) or without (open) LY364947 in HPV− (white) and HPV+ cell lines (red). **(G)** Percentage of annexin V positive cells induced by olaparib on HPV+ SCC47 cells transfected with scramble or anti-miR-182. **(H)** Tumor growth rate of SAS xenografts treated with olaparib, with or without LY2157299, were plotted against days after the first treatment of drugs; control tumor volumes without olaparib treatment were normalized to one; Two-way ANOVA; P < 0.05. Two-tailed Student’s t-test; *, *P* < 0.05; **, *P* < 0.05; ***, *P* < 0.005.

HRR-deficient cells are more sensitive to inhibition of poly(ADP-ribose) polymerase 1 (PARP) due to synthetic lethality, the paradigmatic example of which is that between *BRCA1/2* loss-of-function and PARP inhibition (44). A dose response in the HPV− SAS cell line showed that TGFβ inhibition increased sensitivity to olaparib by 4-fold (Figure 6E). Using annexin-V staining for apoptosis induced by olaparib treated cell lines, we found that HPV+ cell lines were more sensitive to olaparib compared to HPV− cells. However, HPV− cells pretreated with LY364957 were sensitized to olaparib (Figure 6F). SMAD4 knockdown in SAS cells also increased sensitivity to PARP1 inhibition (Supplemental Figure 6, B and C). Moreover, SMAD4 mutant HNSCC cell lines in the Sanger database showed greater sensitivity to olaparib than those in which SMAD4 is wildtype (41) (Supplemental Figure 6D). Importantly, the pronounced sensitivity in HPV+ SCC47 cells to PARP inhibition was rescued by transfection of the miR-182 antagonist (Figure 6G). Olaparib sensitivity is of significant interest clinically and these data suggest a non-responsive cell could be sensitized by inhibiting TGFβ signaling. To test this, we established SAS xenografts and treated them with olaparib with and without TGFβ small molecule inhibitor, LY2157299. As expected, olaparib treatment of TGFβ competent SAS had little effect on tumor growth over the 12-day treatment. However, simultaneous treatment with LY2157299 elicited significant tumor growth inhibition (Figure 6H).

### Loss of TGFβ shifts DNA repair to alt-EJ

Alternative end joining (alt-EJ) pathways (37, 38) are thought to compete, albeit poorly, with HRR for DSB repair in S-phase (39), and back up repair when either HRR or NHEJ is compromised (40). To evaluate the repair from alt-EJ, we established a SAS cell clone with a GFP reporter construct that detects alt-EJ events resulting from I-SceI endonuclease-induced DSB (38). Two different inhibitors of TGFβ signaling significantly increased the frequency of alt-EJ events (Figure 7A). In contrast, a specific PARP1 inhibitor, AG14361, reduced the frequency of alt-EJ events because PARP is necessary for alt-EJ. Thus HNSCC with deficient TGFβ signaling may use alt-EJ more frequently. Indeed, analysis of HNSCC TCGA indicates that HPV+ cancers exhibit significantly greater expression of PARP1 and POLQ, both of which are critical components in alt-EJ (Supplemental Figure 6E).

**Figure 7.**
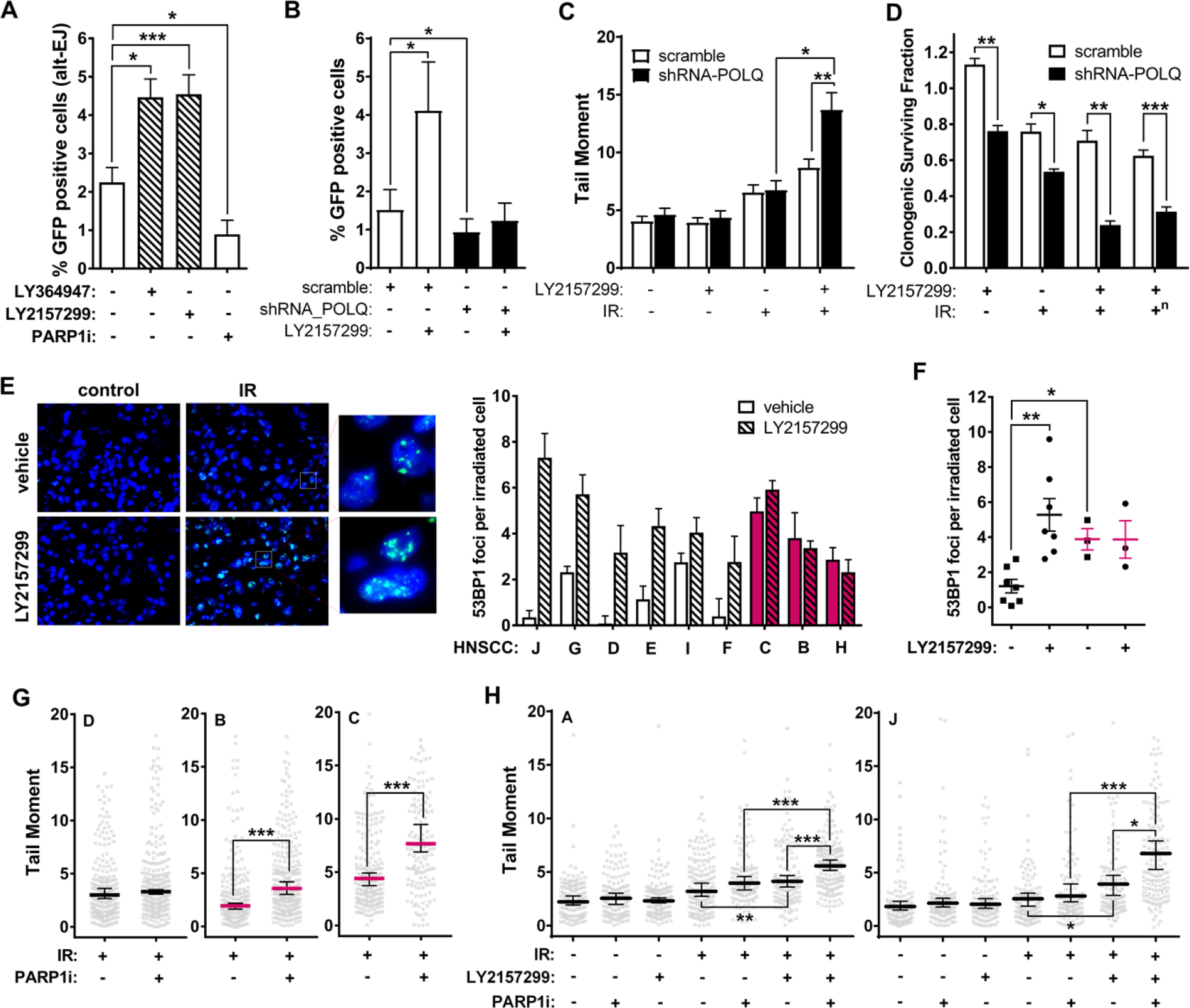
Loss of TGFβ signaling increases reliance on alt-EJ. **(A)** Percentage of GFP-positive cells from I-SceI-alt-EJ reporter cells treated with TβRI inhibitors (LY364947 or LY2157299 at 0.4 ⁊M) or PARP1 inhibator AG14361 (PARP1i) at 1 ⁊M. **(B)** Percentage of GFP-positive cells from I-SceI-alt-EJ reporter cells transfected with shRNA against POLQ or scramble. **(C)** Olive tail moments 5 hr after 0 or 10 Gy irradiation of SAS cells with (scramble) or without POLQ (shRNA-POLQ) treated with LY2157299 or vehicle control for 24 hr. **(D)** Clonogenic surviving fractions of isogenic SAS cell pair transfected with shPOLQ or scrambled RNA exposed to 2 Gy radiation with or without LY2157299. Superscript “n” indicates that difference in LY2157299 alone induced effects has been normalized. **(E)** Representative images of 53BP1 foci in HPV-primary tissue samples from primary HNSCC G (see Supplemental Figure 6H; left panel). Number of 53BP1 foci per cell in irradiated (4 Gy) HPV− (white) and HPV+ (red) HNSCC specimens pretreated with (hatched) or without (open) LY2157299 at 5 hr post irradiation (right panel). **(F)** Data from HNSCC specimens in panel B were grouped according to HPV status and LY2157299 treatments. Panel A to F, two-tailed Student’s t-test; *, *P* < 0.05; **, *P* < 0.05; ***, *P* < 0.005. **(G)** Olive tail moments of irradiated (10 Gy, 5 hr) HPV− (Black) or HPV+ (red) HNSCC (HNSCC_D, B and C) treated with or without AG14361 (i.e. PARP1i). Median values are shown with 95% confidence intervals. **(H)** Olive tail moments of HPV− cells (HNSCC_A and J) treated with or without IR, TβRI inhibitor LY2157299, AG14361, or combinations. Panel G and H, Mann-Whitney U test; *, *P* < 0.05; **, *P* < 0.01; ***, *P* < 0.005.

Alt-EJ is both less efficient and more error-prone than HRR, which leads to more residual damage and cell death. We created POLQ isogenic cell pair in HPV− cell line SAS to investigate roles of POLQ in TGFβ repressed alt-EJ (Supplemental Figure 6F). The low frequency of alt-EJ events in control cells was further reduced in POLQ-deleted cells, and the effect of TGFβ inhibition was completely eliminated (Figure 7B). Alt-EJ promotes genomic instability and contributes to chromosome aberrations, especially translocations (45). Consistent with upregulated alt-EJ after TGFβ inhibition, TGFβ inhibition increased chromosome aberrations (CA) in irradiated HPV− SAS cells but CA were not significantly affected by TGFβ inhibition in cells with POLQ knockdown (Supplemental Figure 6G). The analysis of neutral comet tail-moments reflects DSB that are unrepaired or DNA fragmentation from cells undergoing apoptosis, either of which demonstrates a poor execution of the DDR. Irradiated POLQ deleted cells and control cells displayed similar tail-moments in a neutral comet assay. However when TGFβ signaling was blocked prior to irradiation, tail moment was significantly greater in POLQ deleted cells (Figure 7C). This differential was evident by clonogenic assay, in which survival of POLQ-depleted cells was significantly decreased by TGFβ inhibition or irradiation (Figure 7D). Notably, radiosensitivity is greatest when both TGFβ signaling and POLQ function are inactive. These data led us to conclude cells with defective TGFβ signaling have a specific requirement for POLQ, consistent with a shift to alt-EJ repair.

To validate the relevance of these *in vitro* studies to human HNSCC, we used explants of 10 primary patient HNSCC (Supplemental Figure 6H). Immunostaining of p-SMAD2 indicative of endogenous TGFβ signaling varied in these HNSCC specimens (Supplemental Figure 6I), but the average of HPV+ specimens was significantly lower than that of HPV− specimens (Supplemental Figure 6J). To measure DNA repair proficiency, we used frequency of P53-binding protein 1 (53BP1) foci after irradiation, whose persistence indicate poor DSB repair (46). Explants were treated with and without TGFβ inhibitor LY2157299 prior to irradiation and 53BP1 nuclear foci were measured at 5 hr post-radiation. HPV− HNSCC showed few (~2) 53BP1 foci, consistent with repair proficiency, whereas HPV+ HNSCC exhibited more than twice as many 53BP1 residual foci indicative of poor repair (Figure 7E). As a group, HPV− HNSCC explants in which TGFβ signaling was inhibited showed significantly more persistent 53BP1 foci, which approached the baseline levels of HPV+ explants that were unresponsive TGFβ inhibition (Figure 7F). To confirm this as evidence of unrepaired DNA damage, we employed the neutral comet assay to detect residual DSB in primary HNSCC cancers irradiated as explants. Irradiated HPV+ HNSCC explains showed significantly greater residual DNA damage upon PARP1 inhibition compared to HPV-samples (Figure 7G). We hypothesized that use of TGFβ inhibitors in HPV− specimens would replicate the consequences of HPV infection on PARP sensitivity. Notably, HPV-explants exhibited minimal response to PARP inhibition alone but significantly increased residual DNA damage when combined with LY2157299, also increased unrepaired DSB following exposure to radiation. Moreover, the triple combination of radiation, PARP inhibition and LY2157299 resulted in less repair than either combination (Figure 7H). Thus, loss of TGFβ signaling in HPV+ cancers compromises DNA repair that increases vulnerability to cytotoxic therapy and can be recapitulated in HPV− cancer by pharmaceutical inhibition of TGFβ signaling.

## Discussion

Although the greater sensitivity of HPV+ HNSCC to cytotoxic therapy is well-documented, our studies provide compelling evidence that is the basis for this differential is abrogation of TGFβ signaling. Loss of TGFβ signaling releases suppression of miR-182 that targets both BRCA1 and FOXO3, the latter of is necessary for ATM kinase activity. As both BRCA1 and ATM are essential for HRR, this deficiency increases sensitivity to radiation, cisplatin and PARP inhibition. Unexpectedly, the penultimate mechanism by which HPV sensitizes HNSCC to cytotoxic therapy is a shift to alt-EJ repair, which is more error prone and creates vulnerability by reliance on novel targets, such as POLQ, that can be exploited clinically (47).

The DNA damage response is an underappreciated facet of TGFβ biology (48). The current studies demonstrate that failure of TGFβ signaling from HPV infection or pharmaceutical inhibition redirects DDR pathway choice. Both classical pathways for DSB repair, i.e. NHEJ and HRR, are affected by TGFβ signaling. TGFβ positively regulates ligase IV (15), which is a critical player in NHEJ. The NHEJ repair pathway, which involves direct rejoining of DSB ends, is a fast but error-prone process that is functional throughout the cell cycle. In contrast, the HRR pathway, which uses homologous DNA sequences as a repair template, is a slower, error-free process occurring in S/G2. Our studies show that HRR is specifically decreased upon TGFβ inhibition by either HPV abrogation or pharmaceutical blockade of receptor-mediated TGFβ signaling. We identified TGFβ suppression of miR-182 as the key link between ATM and BRCA1. BRCA1 affects HRR through two major steps: facilitating the processing and resection of DNA broken ends, and binding of RAD51 to ssDNA. ATM is not only needed for initiation but also for completion of HRR (49). Many HRR components are ATM substrates, including BRCA1, BLM, NBS1, MRE11, and CtIP (50). Furthermore, ATM defects are also synthetic lethal with PARP inhibition (51). Notably, ATM phosphorylates BRCA1 at multiple Ser/Thr sites, among which phosphorylation at serine 1423 and serine 1524 are important in precise NHEJ (52). In addition, failure of ATM kinase impairs DSB recognition, which is often assessed by analyzing ⁊H2AX, itself a substrate for ATM phosphorylation (53).

TGFβ inhibition sensitizes brain, breast, and lung cancer cells to radiation-induced clonogenic cell death and improves tumor control in preclinical tumors (20,22,25,28). Blockade of TGFβ signaling also augments the response of brain tumor models to chemoradiation (21,24,54). Teicher and colleagues demonstrated that TGFβ activity contributes to drug resistance and hinders response to chemotherapy in liver and colon cancer preclinical models (55). Pharmacological approaches to block TGFβ signaling include monoclonal antibodies, antisense oligonucleotides, and small molecule inhibitors (56). The consensus of these studies and ongoing trials is that TGFβ inhibition can be achieved safely but the major questions remain as to which patients will benefit and which of its pleiotropic actions contribute to therapeutic outcome.

Abrogation of TGFβ signaling is an extremely frequent event in HNSCC. *SMAD4* deletion in HPV− HNSCC led Wang and colleagues to engineer conditional deletion of *SMAD4* that generated a murine model of spontaneous HNSCC (16). Consistent with our studies showing that loss of TGFβ signaling in HPV+ HNSCC, these HPV-tumors are characterized by genomic instability and decreased HRR (16). HPV infection is functionally equivalent to SMAD4 deletion because oncogenes E5, E6 and E7 essentially abrogate TGFβ signaling. Viral protein E5 decreases phosphorylation of SMAD2 and nuclear translocation of SMAD4, as well as leading to progressive down-regulates T⁊RII (57); E6 renders cells resistant to TGFβ mediated growth control by interacting and degrading the TIP-2/GIPC (58), and E7 interacts constitutively with SMAD2, 3 and 4, which significantly impedes SMAD4-mediated transcriptional activity (59). Hence HPV+ HNSCC provides an ‘experiment of nature’ that demonstrates the critical role of TGFβ signaling in efficient execution of the DNA damage response. Given the significant survival differential between HPV+ and HPV− cancer patients, our studies provide strong rationale to treat cancers in which TGFβ signaling is intact with TGFβ blocking pharmaceuticals with the aim to compromise DNA repair and improve outcome from DNA damaging therapy or PARP inhibitors.

## Authors’ Contributions

Q.L., L.M., I.C. and T.J. performed the experiments; A.L. and M.B. generated tissue microarrays and performed HPV analyses; J.S. and I.C. generated tumor response data; M.H.B-H., J.M., P.H. and Q.L. planned the project; Q.L. and M.H.B-H. prepared the manuscript; L.P., M.A.P. and Q.L. did the bioinformatic analyses. M.H.B.H communicated with the journal, is accountable for fulfilling requests for reagents and resources, and arbitrated decisions and disputes. All authors read and edited the manuscript.

## Acknowledgements

The authors would like to thank Drs. Simon Powell, Henning Willers, George Iliakis and Alba Gonazlez-Junca for helpful comments, DKTK-ROG, and Dr. Jeremy Stark for reagents, and Mr. William Chou and Ms. Shiva Bolourchi for experimental support. This study was funded by the UCSF Department of Radiation Oncology to MHBH.

